# EvoStructCLIP: A Mutation-Centered Multimodal Embedding Model for CAGI7 Variant Effect Prediction

**DOI:** 10.64898/2026.03.02.707336

**Authors:** Kyungkeon Chung, Jaekyung Lee, Yutae Kim, Jeongheon Lee, Junyoung Park, Hyejin Lee

## Abstract

We present EvoStructCLIP, a mutation-centered multimodal embedding model that integrates local 3D structural windows and evolutionary constraints to predict missense variant effects. EvoStructCLIP combines two encoders: a structure voxel encoder derived from AlphaFold residue neighborhoods and an MSA-based evolutionary encoder. It aligns the modalities through CLIP-style contrastive learning, with FuseMix regularization and an auxiliary pathogenicity loss trained on 153,787 ClinVar variants.

Evaluations using lightweight regressors demonstrate that EvoStructCLIP embeddings capture highly transferable predictive signals across diverse phenotypes, including gene-specific functional readouts of *BRCA1, KCNQ4*, and *PTEN*/*TPMT*. This transferability is further supported in the CAGI7 blind competition setting, where models generalized to predicting different gene-specific readouts for *BARD1, FGFR*, and *TSC2* without target-specific retraining and achieved competitive performance across heterogeneous biological tasks.

## 1 Introduction

Despite recent advances in large protein language models and structure prediction frameworks, complete and reliable prediction of thermodynamic stability changes due to mutations remains unsolved [1], despite advances in understanding the molecular structure, stability, and ultimately function [2]. Accurate stability prediction is essential for understanding molecular mechanisms of diseases, and evolution [3]. Although recent deep learning models such as AlphaFold [4], or RoseTTAFold [5], and subsequent models [6, 7] have substantially improved backbone-level structural accuracy, several fundamental challenges persist [8, 9].

One major challenge arises from the intrinsic idiosyncrasy of individual protein molecules. Even within the same fold class or homologous family, subtle sequence variations can induce disproportionately large effects on local packing [10], conformational flexibility [11], or interaction networks [12]. These effects are often mediated by context-specific residue couplings that are not uniformly represented across available training datasets [13]. This uneven representation introduces systematic inductive biases. Models trained predominantly on well-characterized proteins may implicitly encode assumptions that do not generalize whether to all proteins [14] or to other contexts [15, 16]. In such settings, predictive accuracy may reflect familiarity with specific molecular contexts rather than genuine generalization across the protein universe.

Given these constraints, it is reasonable to consider that models tailored to narrower molecular regimes may yield improved performance. Training or fine-tuning strategies conditioned on protein family [17] or structural class [18] may better capture the local regularities governing stability within those regimes. While such specialization may sacrifice broad universality, it acknowledges the heterogeneous nature of protein space and the practical limitations imposed by available data.

Here we present EvoStructCLIP, a small-scale protein window embedding model. We trained the model using a composite supervision strategy using clinically annotated variants, the voxelized representations of structure and the evolutionary constraints around each variant. We show that the pursuit of domain-adaptive or protein-specific modeling frameworks represents a pragmatic complement to large, general-purpose models in multiple downstream tasks. EvoStructCLIP showed good performance in a multitude of tasks at Critical Assessment of Genome Interpretation 7(CAGI7) challenges [19].

## 2 Data preprocessing

### 2.1 Clinically annotated missense variants

ClinVar variants (release July 29, 2025) were used to supervise multimodal representation learning [20]. The dataset was further restricted to single-nucleotide missense substitutions only. Variants annotated as pathogenic or likely pathogenic were assigned to the pathogenic class, whereas variants annotated as benign or likely benign were assigned to the benign class. Variants with conflicting interpretations or uncertain clinical significance were excluded.

Variants were mapped to canonical UniProtKB isoforms to ensure residue indexing consistency [21]. Sequence alignment against Swiss-Prot and TrEMBL (UniProt release 2025_03) [21] confirmed reference concordance. Variants failing structural mapping or isoform consistency checks were excluded. Exact duplicates were removed after transcript-protein identifier matching.

The final dataset comprised 153,787 unique missense variants with high-confidence binary pathogenicity annotations.

### 2.2 Voxel representations of the local tertiary structure

To represent the local tertiary environment around each mutated residue, we constructed three-dimensional voxel representations from AlphaFold DB human proteome protein models (release v4; UP000005640) [22]. For each variant, a grid of size 7 × 7 × 7 with a voxel spacing of 2 Å was centered on the C*α* atom of the mutated residue.

Each voxel was annotated with 42 closeness channels corresponding to the proximity of C*α* and C*β* atoms for each of the 21 amino acid types, following the general strategy of Gao *et al*. [23]. Closeness values were computed as a normalized linear decay of the minimum atomic distance within a 5 Å radius.

Additional structural descriptors were incorporated to enrich the representation. Relative sequence position was encoded using normalized signed and absolute sequence distances between the mutated residue and the residue mapped to each voxel based on nearest C*α* coordinates. Per-residue AlphaFold confidence scores (pLDDT) were projected onto the grid to reflect structural reliability [24]. Local dynamic flexibility was quantified using mean square fluctuations derived from the Gaussian network model (GNM) analysis implemented in ProDy [25].

Together, these components yielded a 46-channel voxel representation integrating geometric proximity, positional context, model confidence, and structural dynamics.

### 2.3 Evolutionary constraints of the local primary structure

Evolutionary constraints were captured using multiple sequence alignments (MSAs) generated with MMseqs2 [26] against the UniRef90 database (UniProt release 2025_03) [21]. Sequence searches were performed with an E-value threshold of 1 × 10^−3^ and a maximum of 500 retrieved sequences per query.

To ensure alignment quality and control redundancy, the resulting MSAs were filtered according to the following criteria: (i) maximum pairwise sequence identity of 95%, (ii) minimum sequence identity to the query of 30%, and (iii) minimum alignment coverage of 30%. These thresholds were applied to retain homologous sequences while reducing excessive redundancy and poorly aligned fragments.

### 2.4 Downstream Tasks

#### 2.4.1 *BRCA1* functional and RNA score prediction

To evaluate embedding utility, we regressed on two experimentally calculated values from Findlay *et al*. [27] (urn:mavedb:00000097) downloaded from MaveDB [28] as a downstream task. The predicted values were the functional score, which represents the change in variant frequency under selection reflecting *BRCA1* essentiality, and the RNA score which represents relative mRNA abundance derived from targeted RNA sequencing.

#### 2.4.2 Prediction of normalized *KCNQ4* channel activity

The second downstream evaluation task involved predicting the normalized current *KCNQ4* activity score for each missense variant, as reported by Zheng *et al*. [29] (urn:mavedb:00000094-a-2). This score quantifies the relative channel current produced by each variant compared to the homozygous wild-type baseline and was obtained from MaveDB [28].

#### 2.4.3 *PTEN* and *TPMT* VAMP-seq abundance prediction

For the third downstream evaluation, we used variant-level abundance measurements for *PTEN* (urn:mavedb:00000013-a-1) and *TPMT* (urn:mavedb:00000013-b-1) obtained from MaveDB [28]. These data were generated using the VAMP-seq assay [30], which quantifies steady-state intracellular protein abundance as a proxy for variant stability. The regression target was a continuous abundance score corresponding to the relative protein level of each missense variant compared to the wild-type reference.

#### 2.4.4 Ablation analysis with random embedding

To show that our embedding is meaningful, for each of the downstream tasks, we replaced the embedding from EvoStructCLIP with a randomized vector with the same dimension for comparison.

## 3 Model & training

### 3.1 Model architecture

EvoStructCLIP is a multimodal framework designed to jointly model residue-level structural context and evolutionary constraint. The architecture is motivated by the hypothesis that local three-dimensional packing effects and sequence conservation encode complementary signals of mutational impact. To capture these modalities, the model comprises (i) a voxel-based encoder that represents the spatial microenvironment surrounding the mutated residue and (ii) an MSA-based encoder that captures conservation and substitution patterns across homologous sequences. The two embedding spaces are aligned through a shared optimization objective, encouraging consistent representations across structural and evolutionary views of the same variant. (Figure 1).

**Figure 1.**
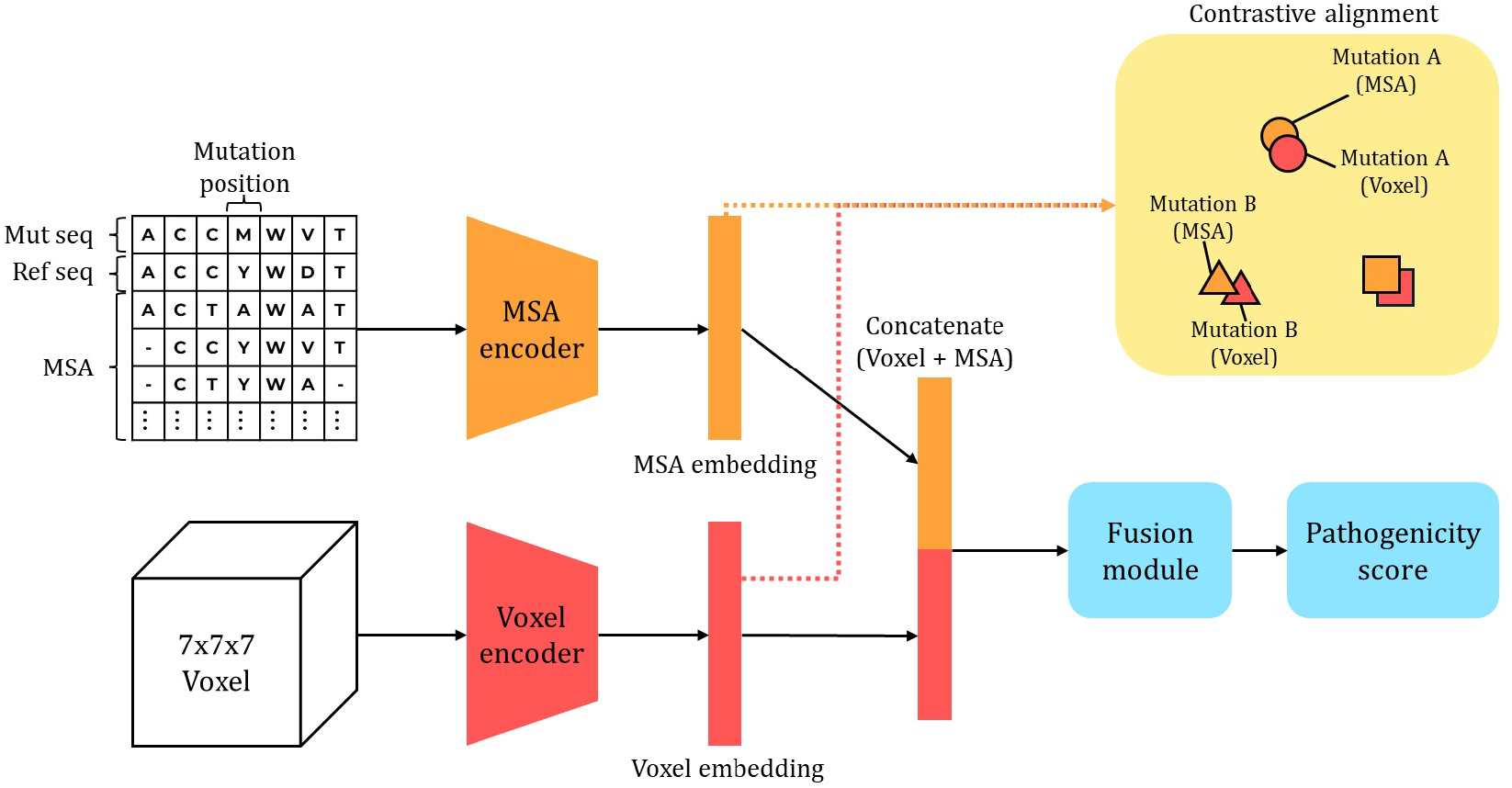
Overview of the EvoStructCLIP framework. Structural inputs are converted into voxel grids centered on the mutated residue and processed by the voxel encoder to produce structural embeddings. In parallel, multiple sequence alignments are processed by the MSA encoder to produce evolutionary embeddings. The embeddings are jointly optimized using (i) a supervised classification loss for pathogenicity prediction and (ii) a CLIP-style contrastive loss computed from a cross-modal similarity matrix to enforce alignment between structural and evolutionary representations.

#### 3.1.1 Voxel encoder

The voxel encoder processes the 3D voxelized structural features described earlier. The backbone consists of stacked 3D MBConv blocks with pointwise expansion, depthwise convolution, and squeeze-and-excitation attention [31, 32], following the efficiency principles of EfficientNet [33]. The output feature maps are then refined by a three-dimensional coordinate attention module (CoordAtt3D) [34], which captures long-range dependencies along the depth, height, and width axes before global pooling.

Following global average pooling, the resulting structural representation is augmented with mutation-specific information. The wild-type and substituted residues are embedded independently, concatenated, and transformed through a non-linear projection to yield a compact mutation descriptor. This descriptor is integrated with the pooled structural vector, thereby explicitly encoding both the local three-dimensional context and the biochemical identity of the residue substitution. A final lightweight projection layer produces the voxel-derived embedding used in downstream multimodal alignment and classification.

#### 3.1.2 MSA encoder

The MSA encoder models evolutionary information from sequence alignments centered on the mutated position. Each amino acid is embedded into a continuous representation, yielding a tensor of shape (*L, D, C*), where *L* denotes the alignment window length, *D* the MSA depth, and *C* the embedding dimension.

To capture complementary dependencies along both axes of the alignment, we introduce a cross-axial Mamba block (Figure 2). Along the sequence length axis, a state-space layer enables efficient long-range context propagation [35, 36], while along the alignment depth axis, localized convolutional filters extract consensus patterns across homologous sequences at the mutation site. This design allows the encoder to jointly model positional conservation and depth-wise evolutionary variability in a structured manner [37]. Residual connections and RMS normalization are used to stabilize training [38].

**Figure 2.**
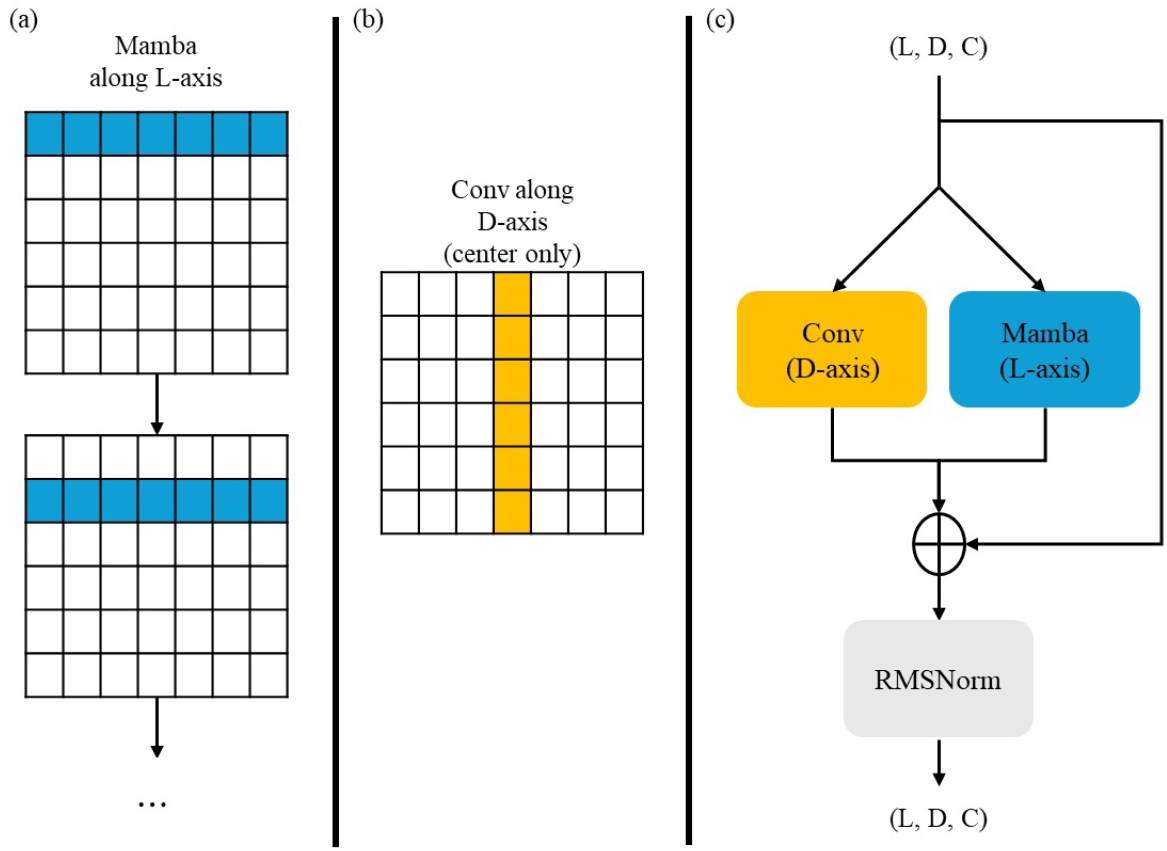
Cross-axial Mamba block used in the MSA encoder. (a) A Mamba state-space layer is applied along the sequence length axis (*L*), operating independently across depth positions. (b) A 1D convolution is applied along the alignment depth axis (*D*) at the central sequence position corresponding to the mutated residue. (c) The representations from both directions are integrated by residual addition, followed by RMSNorm, resulting in an updated tensor of shape (*L, D, C*). Per-axis normalization is applied before each operation (not shown).

### 3.2 Objective Functions

EvoStructCLIP is trained end-to-end using a composite objective function that simultaneously optimizes task performance and aligns the latent representations of different modalities. The total objective function ℒ_*total*_ is formulated as a weighted sum of three distinct loss components: a variant pathogenicity classification loss (ℒ_*cls*_), a cross-modal contrastive loss (ℒ_*clip*_), and a latent augmentation loss (ℒ_*fusemix*_).

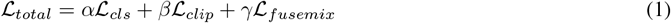

where *α, β, γ* are hyperparameter coefficients controlling the contribution of different loss components.

#### 3.2.1 Pathogenicity Loss

To learn robust multimodal representations that generalize across various downstream tasks, the model predicts ClinVar missense pathogenicity as one of the objectives.

To integrate the variant pathogenicity directly into the computation of loss, we append a lightweight feed-forward network to the model. Specifically, the concatenated voxel and MSA embeddings are passed through a linear projection, batch normalization, SiLU activation, and a final linear layer. This network produces a scalar pathogenicity score, which serves as the output logit *z*_*i*_ for the *i*-th variant. For a batch of *N* samples, we apply the standard binary cross-entropy (BCE) loss between the true clinical labels and the predicted probabilities. Let *y*_*i*_ ∈ {0, 1} be the ground-truth pathogenicity label, and *ŷ*_*i*_ = *σ*(*z*_*i*_) be the predicted probability obtained by applying the sigmoid function *σ*(·) to the logit *z*_*i*_. The classification loss is defined as:

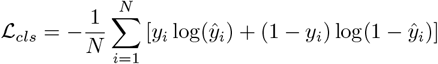

#### 3.2.2 CLIP Loss

To align the structural and evolutionary features in the latent space, we employ a symmetric contrastive loss [39]. Let **v**_*i*_ and **m**_*i*_ denote the *L*_2_-normalized feature embeddings extracted from the voxel and MSA encoders, respectively, for the *i*-th sample in a batch of size *N* .

The scaled pairwise similarity between the *i*-th voxel and the *j*-th MSA embedding is defined as 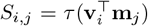, where *τ* is a learnable temperature parameter. Using these similarity scores, the voxel-to-MSA and MSA-to-voxel contrastive losses are computed as:

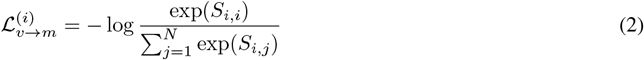

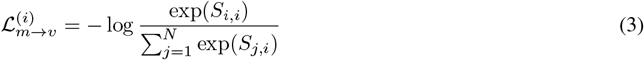

The final CLIP loss is the average of these two symmetric components across all samples:

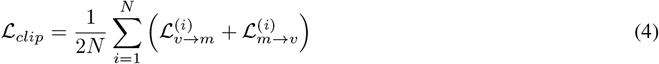

#### 3.2.3 FuseMix Loss

To further regularize the feature space and improve the model’s robustness against data scarcity, we employ FuseMix [40] as an auxiliary latent-space augmentation strategy, which builds upon the foundational mixup technique [41]. While the original framework utilized this approach as a primary alignment objective for frozen encoders, we adapt it as a supplementary regularization term to stabilize the end-to-end training of our architecture. As proposed by Vouitsis *et al*. [40], the interpolation is systematically applied to the unnormalized embeddings **h**_*v*_ and **h**_*m*_ prior to *L*_2_ normalization.

For any two distinct samples *i* and *j* drawn from a given batch, we sample a shared mixing coefficient *λ* from a Beta distribution, *λ* ∼ Beta(*α, α*). The mixed raw embeddings are computed as convex combinations:

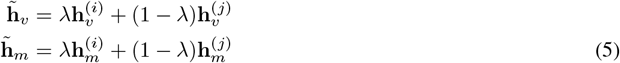

Applying the same mixing coefficient *λ* to both modalities preserves cross-modal pairing between the structural and evolutionary representations during interpolation. These interpolated vectors are subsequently *L*_2_-normalized to yield the augmented representations: 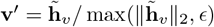 and 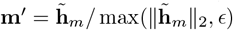.

Finally, the symmetric contrastive loss (analogous to Equation 4) is calculated for these mixed, normalized embeddings to compute the auxiliary loss, ℒ_*fusemix*_. By incorporating this term alongside the primary classification and original CLIP objectives, we effectively encourage the encoders to learn a smoother, more generalized latent space without compromising the integrity of the original pathogenic signals.

### 3.3 Training scheme

Optimization was carried out using AdamW [42] with cosine annealing learning rate scheduling [43] for a maximum of 100 epochs. Model selection was guided by validation precision–recall area under the curve (PR-AUC) [44]. Data augmentation comprised random voxel rotations and reflections [45] and stochastic shuffling of MSA depth.

### 3.4 Missense Variant Effect Evaluation

To assess whether pathogenicity information was effectively encoded in the embeddings, we held out 10% of ClinVar variants during training for evaluation. The same classification head architecture used in training was applied at test time.

### 3.5 Downstream evaluation tasks

For each of the gene-specific regression tasks of *BRCA1, KCNQ4*, and *PTEN*/*TPMT*, multimodal embeddings of missense variants from EvoStructCLIP are concatenated with a set of residue- and mutation-level descriptors derived from structural and evolutionary statistics and used to train non-neural network regression models, specifically random forest (RF) [46] and XGBoost (XGB) [47], to predict dataset-specific quantitative phenotypes.

The 19-dimensional handcrafted features concatenated include: pLDDT confidence scores at the mutated residue and neighboring positions, mean square fluctuations from Gaussian network model (GNM) analysis [25], long-range contact order [48], physicochemical property changes upon mutation (volume, hydropathy, and charge differences) [49], predicted folding stability change (ΔΔ*G*) from EvoEF2 [50], backbone torsion angles represented as sine and cosine terms, binary indicators for glycine or proline substitutions, and evolutionary conservation features such as ΔPSIC [51] and Shannon entropy [52].

For all downstream evaluation tasks, we adopted a consistent evaluation protocol. Variants were first partitioned into training and test sets using a 95%/5% split, with 5% of variants held out as an independent test set prior to any model training or hyperparameter optimization. The remaining 95% of the data was used exclusively for model development, including hyperparameter selection and performance estimation via 10-fold cross-validation. Final model performance was assessed on the held-out 5% test set to provide an unbiased estimate of generalization. Additionally, to assess the contribution of the learned multimodal representation in the downstream tasks, we performed an ablation experiment in which the pretrained EvoStructCLIP embedding was replaced with a randomly initialized 256-dimensional vector of identical dimensionality. The handcrafted structural and evolutionary descriptors, regression models (RF and XGB), data splits, and cross-validation protocol were kept unchanged for the evaluation.

## 4 Results

### 4.1 Variant effect prediction on ClinVar validation set

EvoStructCLIP achieved a PR-AUC of 0.926, receiver operating characteristic area under the curve (ROC-AUC) of 0.953, and accuracy of 0.904 on the held-out validation set (Table 1), indicating high discriminative performance in distinguishing pathogenic from benign missense variants. Notably, the MSA-only encoder maintained high performance (PR-AUC = 0.911, ROC-AUC = 0.943, accuracy = 0.895), suggesting that the contrastive alignment enables evolutionary embeddings to internalize structural signals even without explicit structural inputs. The small performance gap suggests that contrastive alignment enables evolutionary embeddings to internalize structural signal. These results show that EvoStructCLIP encodes clinically relevant mutational signals and that contrastive alignment yields robust cross-modal transfer.

**Table 1:** Variant effect prediction performance on the ClinVar validation set.

| Model | PR-AUC | ROC-AUC | Accuracy |
| --- | --- | --- | --- |
| EvoStructCLIP (MSA + Structure) | 0.926 | 0.953 | 0.904 |
| MSA encoder only | 0.911 | 0.943 | 0.895 |

### 4.2 Prediction of *BRCA1* functional and RNA scores

For the *BRCA1* functional score prediction task, both RF and XGB models trained on EvoStructCLIP embeddings achieved strong predictive performance (Table 2). The RF ensemble attained a Pearson correlation of 0.760 with an RMSE of 0.711, while XGB further improved performance to Pearson *r* = 0.789 and RMSE = 0.653. Performance remained stable across folds (mean Pearson *r* = 0.756 *±* 0.012 for RF and 0.782 *±* 0.012 for XGB), indicating consistent generalization across training splits. In contrast, substituting the pretrained embedding with random vectors resulted in substantial performance degradation for both targets, demonstrating that pretrained embeddings provide substantial predictive information beyond handcrafted features.

**Table 2:** Performance on the held-out *BRCA1* test set.

| Target | Model | Pearson $r$ | Spearman $\rho$ | RMSE |
| --- | --- | --- | --- | --- |
| **Functional score** |  |  |  |  |
|  | RF (EvoStructCLIP, ensemble) | 0.760 | 0.634 | 0.711 |
| | RF (EvoStructCLIP, fold test) | $0.756 \pm 0.012$ | $0.630 \pm 0.011$ | $0.712 \pm 0.009$ |
|  | RF (Random embedding, ensemble) | 0.687 | 0.610 | 0.825 |
| | RF (Random embedding, fold test) | $0.682 \pm 0.012$ | $0.607 \pm 0.014$ | $0.826 \pm 0.007$ |
|  | XGBoost (EvoStructCLIP, ensemble) | **0.789** | **0.681** | **0.653** |
| | XGBoost (EvoStructCLIP, fold test) | $0.782 \pm 0.012$ | $0.672 \pm 0.013$ | $0.659 \pm 0.010$ |
|  | XGBoost (Random embedding, ensemble) | 0.699 | 0.630 | 0.754 |
| | XGBoost (Random embedding, fold test) | $0.691 \pm 0.008$ | $0.620 \pm 0.018$ | $0.757 \pm 0.008$ |
| **RNA score** |  |  |  |  |
|  | RF (EvoStructCLIP, ensemble) | **0.612** | **0.335** | **0.571** |
| | RF (EvoStructCLIP, fold test) | $0.599 \pm 0.022$ | $0.320 \pm 0.031$ | $0.574 \pm 0.009$ |
|  | RF (Random embedding, ensemble) | 0.436 | 0.228 | 0.653 |
| | RF (Random embedding, fold test) | $0.401 \pm 0.073$ | $0.205 \pm 0.040$ | $0.654 \pm 0.009$ |
|  | XGBoost (EvoStructCLIP, ensemble) | 0.603 | 0.266 | 0.568 |
| | XGBoost (EvoStructCLIP, fold test) | $0.592 \pm 0.021$ | $0.263 \pm 0.023$ | $0.571 \pm 0.007$ |
|  | XGBoost (Random embedding, ensemble) | 0.531 | 0.280 | 0.630 |
| | XGBoost (Random embedding, fold test) | $0.507 \pm 0.024$ | $0.248 \pm 0.039$ | $0.631 \pm 0.007$ |

*BRCA1* RNA abundance score predictions showed moderate but consistent performance. The RF ensemble achieved Pearson *r* = 0.612 and RMSE = 0.571, while XGB achieved Pearson *r* = 0.603 with comparable RMSE. Cross-validation stability remained consistent (mean Pearson *r* = 0.599 *±* 0.022 for RF and 0.592 *±* 0.021 for XGB). As above, random-embedding baselines showed performance degradation, suggesting that EvoStructCLIP captures aspects of both expression-level consequences and functional impairment.

For CAGI7’s RNA abundance and cell survival prediction of *BARD1* challenge, the model trained to predict functional scores with *BRCA1* data was used for inference and submitted.

### 4.3 Prediction of *KCNQ4* electrophysiological current

In the KCNQ4 normalized-current task, EvoStructCLIP-based RF ensemble achieved a Pearson correlation of 0.568 and Spearman correlation of 0.431, with an RMSE of 0.389 (Table 3). Cross-validation stability was maintained (mean Pearson *r* = 0.563 *±* 0.020), indicating consistent generalization across folds. The XGB ensemble showed comparable performance (Pearson *r* = 0.553, RMSE = 0.394) with similar stability (mean Pearson *r* = 0.536 *±* 0.019). Performance was lower than for the *BRCA1* dataset, consistent with the greater biophysical complexity of electrophysiological current phenotypes. Replacing the pretrained EvoStructCLIP embeddings with random embeddings resulted in a consistent reduction in performance in the task, indicating that the learned multimodal representation contributes substantially to the biophysical phenotype prediction.

**Table 3:** Performance on the held-out *KCNQ4* normalized current test set.

| Target | Model | Pearson $r$ | Spearman $\rho$ | RMSE |
| --- | --- | --- | --- | --- |
| **Normalized current** |  |  |  |  |
|  | RF (EvoStructCLIP, ensemble) | **0.568** | 0.431 | **0.389** |
| | RF (EvoStructCLIP, fold test) | $0.563 \pm 0.020$ | $0.429 \pm 0.014$ | $0.391 \pm 0.007$ |
|  | RF (Random embedding, ensemble) | 0.417 | 0.379 | 0.429 |
| | RF (Random embedding, fold test) | $0.414 \pm 0.010$ | $0.375 \pm 0.011$ | $0.430 \pm 0.002$ |
|  | XGBoost (EvoStructCLIP, ensemble) | 0.553 | **0.438** | 0.394 |
| | XGBoost (EvoStructCLIP, fold test) | $0.536 \pm 0.020$ | $0.426 \pm 0.023$ | $0.400 \pm 0.007$ |
|  | XGBoost (Random embedding, ensemble) | 0.419 | 0.348 | 0.429 |
| | XGBoost (Random embedding, fold test) | $0.400 \pm 0.028$ | $0.335 \pm 0.026$ | $0.435 \pm 0.007$ |

For CAGI7’s gain-of-function variant prediction challenge for *FGFR*, the model trained with *KCNQ4* data was used for inference and submitted.

### 4.4 Prediction of *PTEN/TPMT* abundance

For the *PTEN/TPMT* abundance prediction task, the 10-fold RF ensemble achieved a Pearson correlation of 0.734, Spearman correlation of 0.695, and RMSE of 0.239 (Table 4). Performance across folds was highly stable (mean Pearson *r* = 0.731 *±* 0.003), indicating minimal sensitivity to training partitioning. The 10-fold XGB ensemble demonstrated comparable correlation performance (Pearson *r* = 0.736, Spearman *ρ* = 0.685) with a slightly lower RMSE of 0.233 and similar cross-validation stability (mean Pearson *r* = 0.732 *±* 0.005). While both models achieved nearly identical correlation metrics, XGB exhibited modestly reduced prediction error, suggesting improved modeling of abundance variation.

**Table 4:**
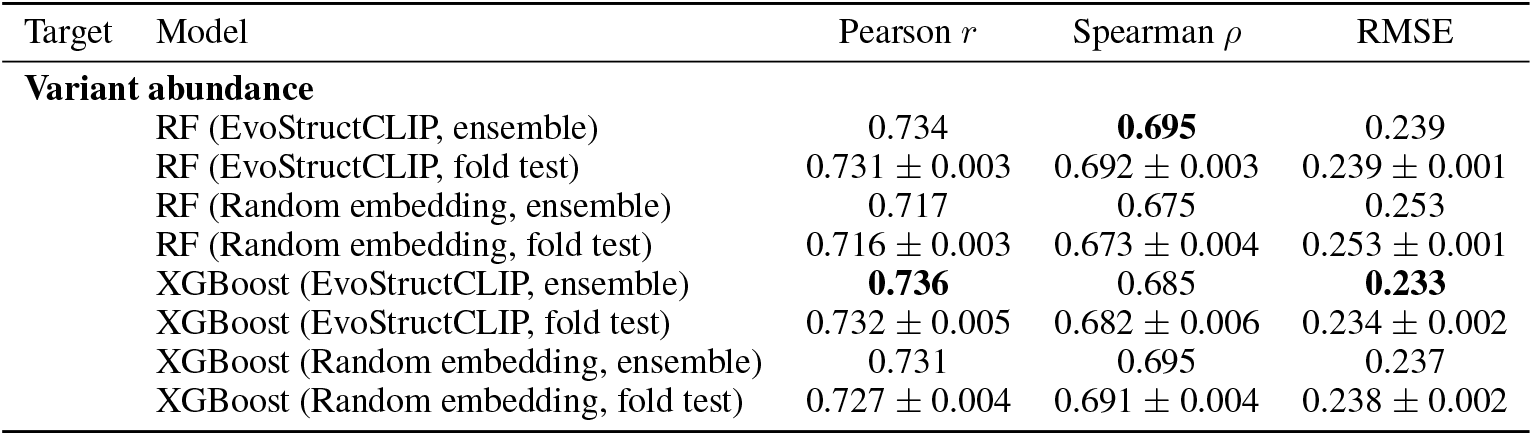
Performance on the held-out *PTEN/TPMT* test set.

In contrast to the *BRCA1* and *KCNQ4* tasks, replacing the pretrained EvoStructCLIP embedding with randomly initialized vectors resulted in only a modest performance decrease (Table 4). These results suggest that handcrafted descriptors account for a substantial fraction of the predictive signal, with multimodal embeddings providing incremental improvement. However, the EvoStructCLIP embedding consistently provided small but measurable improvements in RMSE, suggesting that the learned representation contributes complementary information about local mutation effects. For *TSC2* protein stability prediction challenge, the functional score model trained on *PTEN*/*TPMT* data was used for inference and submitted.

## 5 Conclusion

We introduce EvoStructCLIP, a mutation-centered multimodal embedding framework that integrates local 3D structural context and evolutionary constraints. By aligning voxel-derived structural representations with MSA-derived evolutionary embeddings through a CLIP-style contrastive objective [39] augmented with FuseMix [40] and a pathogenicity loss, the model captures transferable variant-level signals.

Motivated by the intrinsic heterogeneity of protein space, EvoStructCLIP adopts a strategy of creating a generalized embedding followed by fine-tuning on individual proteins rather than relying exclusively on global representations. By constraining learning to mutation-centered structural windows and corresponding evolutionary neighborhoods, the model emphasizes context-specific residue interactions that are often diluted in large-scale, protein-wide embeddings. The contrastive alignment between structural voxels and evolutionary profiles further enforces a shared representation space that captures mechanistic regularities at the mutation level.

Across multiple evaluation settings in the CAGI challenge, EvoStructCLIP demonstrated competitive performance in diverse molecular prediction tasks. Because CAGI is organized as a community-wide, blind competition, all submissions were generated without access to held-out ground truth labels. For the CAGI7 gain-of-function variant prediction challenge for *FGFR*, the model trained on *KCNQ4* functional data was used directly for inference and submitted without target-specific retraining. For the *TSC2* protein stability prediction challenge, the functional score model trained on combined *PTEN*/*TPMT* abundance data was applied for inference and submission. For the CAGI7 RNA abundance and cell survival prediction challenge for *BARD1*, the model trained to predict functional scores using *BRCA1* data was used for inference.

Despite the absence of task-specific fine-tuning, the model achieved strong results in thermodynamic stability prediction in *TSC2*, in modeling RNA abundance and survival effects for *BARD1*, and in predicting activation probability for *FGFR*. These tasks span protein stability, transcript-level abundance, and receptor activation, each dependent on accurately modeling localized structural and evolutionary perturbations. The consistent performance across these heterogeneous endpoints indicates that multimodal alignment at the mutation scale captures transferable mechanistic signal across proteins, assays, and phenotypic readouts.

Taken together, these findings support a complementary paradigm to large, general-purpose protein models. Rather than assuming uniform inductive structure across the protein universe, EvoStructCLIP explicitly models gene-mutation-centered contexts and leverages composite supervision from clinical annotations, structural geometry, and evolutionary variation. While such specialization may not aim for universal coverage, it offers a pragmatic and mechanistically grounded strategy for variant effect prediction under realistic data constraints. Domain-adaptive, protein-focused architectures may therefore serve not as substitutes for foundation-scale models, but as targeted frameworks for extracting stability and functional signals within heterogeneous molecular regimes. Our findings suggest that mutation-centered multimodal embeddings can transfer across diverse protein contexts, offering a practical complement to foundation models.

## 6 Code Availability

Code and training scripts are available at: https://github.com/uxfacdev/CAGI.

